# Noradrenergic circuits in the forebrain control affective responses to novelty

**DOI:** 10.1101/2020.04.11.037200

**Authors:** Daniel Lustberg, Rachel P. Tillage, Yu Bai, Molly Pruitt, L. Cameron Liles, David Weinshenker

## Abstract

**Rationale:** In rodents, exposure to novel environments elicits initial anxiety-like behavior (neophobia) followed by intense exploration (neophilia) that gradually subsides as the environment becomes familiar. Thus, innate novelty-induced behaviors are useful indices of anxiety and motivation in animal models of psychiatric disease. Noradrenergic neurons are activated by novelty and implicated in exploratory and anxiety-like responses, but the role of norepinephrine (NE) in neophobia has not been clearly delineated.

**Objective:** We sought to define the role of central NE transmission in neophilic and neophobic behaviors.

**Methods:** We assessed dopamine β-hydroxylase knockout (*Dbh -/-*) mice lacking NE and their NE-competent (*Dbh +/-*) littermate controls in neophilic (novelty-induced locomotion; NIL) and neophobic (novelty-suppressed feeding; NSF) behavioral tests with subsequent quantification of brain-wide c-fos induction. We complimented the gene knockout approach with pharmacological interventions.

**Results:** *Dbh -/-* mice exhibited blunted locomotor responses in the NIL task and completely lacked neophobia in the NSF test. Neophobia was rescued in *Dbh -/-* mice by acute pharmacological restoration of central NE with the synthetic precursor L-3,4-dihydroxyphenylserine (DOPS), and attenuated in control mice by the inhibitory α2-adrenergic autoreceptor agonist guanfacine. Following either NSF or NIL, *Dbh -/-* mice demonstrated reduced c-fos in the anterior cingulate cortex, medial septum, ventral hippocampus, bed nucleus of the stria terminalis, and basolateral amygdala.

**Conclusion:** These findings indicate that central NE signaling is required for the expression of both neophilic and neophobic behaviors. Further, we describe a putative noradrenergic novelty network as a potential therapeutic target for treating anxiety and substance abuse disorders.

## Introduction

A novel environment may represent either an appetitive opportunity or a potential threat; the valence of the context is uncertain because it has never been encountered before (Kafkas and Montaldi 2018). In novel environments, it is adaptive for animals to exhibit initial anxiety-like behavior (neophobia) that gradually transitions to exploratory behavior (neophilia) once the environment has proven to be non-threatening (Barnett 1958; Dulawa et al. 2005; Osorio-Gómez et al. 2018). Novelty-seeking and novelty-fearing personality traits correlate with substance abuse and anxiety disorders, respectively (Bardo et al. 1996; Dovey et al. 2008; Marcontell et al. 2003; Tulving et al. 1994; Weierich et al. 2010). In animal models of psychiatric disease, behavioral responses to novel environments are also strong predictors of drug-seeking and anxiety-like behavior (Dulawa 2009; Pawlak et al. 2008; Walker et al. 2009; Wingo et al. 2016).

Neophilic behavior can be assessed in mice with the novelty-induced locomotion (NIL) test, which measures exploration in a novel environment (Kabbaj et al. 2000; Walker et al. 2009). In the NIL test, high locomotor reactivity to novelty is interpreted as increased exploratory behavior, while low locomotor responses are interpreted as decreased exploratory behavior (Cubells et al. 2016; Schroeder et al. 2013; Stone et al. 1999). Neophobia can be assessed in mice with the novelty-suppressed feeding (NSF) test, a conflict-based model of anxiety-like behavior in which anxiety elicited by exposure to an unfamiliar environment suppresses feeding in hungry mice presented with food (Cryan and Sweeney 2011; Dulawa et al. 2005). In the NSF test, longer latencies to begin feeding are interpreted as increases in neophobia.

Novel environments activate neural circuits involved in spatial memory (Takeuchi et al. 2016; Wagatsuma et al. 2018), attention (Aston-Jones et al. 2007; Gompf et al. 2010), motivation (Rinaldi et al. 2010; Struthers et al. 2005), arousal (Britton and Indyk 1990; Stone et al. 2005), and anxiety (Delini-Stula et al. 1984b; File 2001; Sheth et al. 2008), which enables flexible responding to potential threats or rewards that may emerge under uncertain conditions (Aston-Jones et al. 1999; Kafkas and Montaldi 2018; Snyder et al. 2012). Although much is known about the biological mechanisms that drive conditioned responses to familiar contexts in animal models of anxiety and drug addiction (Koob and Simon 2009; Koob and Volkow 2016; Martin et al. 2009; Tovote et al. 2015), the basic neurochemistry and circuitry of the network that controls innate behavioral responses to novel environments remain largely unknown.

The locus coeruleus (LC) is the primary source of norepinephrine (NE) to the forebrain, projecting extensively within circuits that govern attention, arousal, emotion, and motivated behavior (Aston-Jones et al. 1999; Berridge et al. 1996; Sara and Bouret 2012; Uematsu et al. 2015). A longstanding theory is that the LC-NE system is sensitive to contextual novelty (Gompf et al. 2010; Sara et al. 1995), and it is well established that the firing rate of LC neurons increases in novel environments and declines as the context becomes familiar (Aston-Jones and Bloom 1981; Vankov et al. 1995).

Hyperactivity of the LC-NE system has been implicated in the pathophysiology of anxiety and substance abuse disorders (Aston-Jones et al. 1994; Brady 1994; Goddard et al. 2010; Tanaka et al. 2000; Weinshenker and Schroeder 2007; West et al. 2009), and drugs that suppress the activity of the LC or reduce NE signaling have therapeutic efficacy for patients with these conditions (Aston-Jones and Kalivas 2008; Boehnlein and Kinzie 2007; Fox et al. 2012). Experimental disruptions of the LC-NE system also suppress innate behavioral responses to novel environments and stimuli (Archer et al. 1981; Delini-Stula et al. 1984a; Harro et al. 1995; Neophytou et al. 2001; Sara et al. 1995; Stone et al. 1999).

The LC-NE system innervates a constellation of cortical and limbic regions implicated in spatial memory and novelty detection, including the anterior cingulate cortex (ACC), medial septum/diagonal band (MS/DB) complex, and hippocampus (Chandler 2016; Guiard et al. 2008; Uematsu et al. 2015). LC-NE signaling within the basolateral amygdala (BLA) elicits anxiety-like behavior (McCall et al. 2017; Siuda et al. 2016), and NE transmission within the bed nucleus of the stria terminalis (BNST) is associated with stress responses, innate fear, and reward seeking (Avery et al. 2016; Lebow and Chen 2016; Weinshenker and Schroeder 2007). LC-NE inputs to the anterior insula (AI) and medial orbitofrontal cortex (MO) modulate decision making, emotion, and motivated behavior (Mather et al. 2016; Petrides 2007; Rojas et al. 2015; Sadacca et al. 2017). Thus, the LC is well poised to orchestrate both exploratory and anxious behavior in unfamiliar environments (Berridge and Dunn 1989; McCall et al. 2015; Snyder et al. 2012).

In the present study, we investigated the consequences of genetic or pharmacological disruption of central NE signaling on neophilic and neophobic behavior in the NIL and NSF tests using NE-deficient (dopamine β-hydroxylase knockout; *Dbh -/-*) mice and NE-sufficient (*Dbh +/-*) controls (Thomas et al. 1995). We also determined the effects of pharmacological restoration of central NE synthesis on neophobia in *Dbh -/-* mice and pharmacological reduction of NE transmission on neophobia in control mice (Mineur et al. 2015; Thomas et al. 1998). Finally, we measured how genetic NE deficiency alters c-fos induction in the LC and 15 of its target structures in the forebrain as a proxy measure of task-specific neuronal activity following NIL and NSF.

## Methods

### Subjects

*Dbh -/-* mice were maintained on a mixed 129/SvEv and C57BL/6J background, as described (Thomas et al. 1998; Thomas et al. 1995). Pregnant *Dbh +/-* dams were given drinking water supplemented with the β-adrenergic receptor (βAR) agonist isoproterenol and α1AR agonist phenylephrine (20 μg/ml each) + vitamin C (2 mg/ml) from E9.5-E14.5, and L-3,4-dihydroxyphenylserine (DOPS; 2 mg/ml + vitamin C 2 mg/ml) from E14.5-parturition to prevent developmental lethality associated with homozygous *Dbh* deficiency (Mitchell et al. 2008; Thomas et al. 1995). *Dbh -/-* mice are easily identified by their ptosis phenotype, and genotypes were confirmed by PCR, as described (Mitchell et al. 2008; Thomas et al. 1995). *Dbh +/-* mice behavior and NE levels are indistinguishable from wild-type (*Dbh +/+*) mice and were used as controls (Bourdélat-Parks et al. 2005; Mitchell et al. 2006).

All experiments included adult mice (3-8 months old) of both sexes. Because sex differences were not reported in the literature nor observed in pilot experiments, male and female mice of the same *Dbh* genotype were pooled. All animal procedures and protocols were congruent with the National Institutes of Health guidelines for the care and use of laboratory animals and were approved by the Emory University Animal Care and Use Committee. Mice were maintained on a 12 h light/12 h dark cycle with access to food and water *ad libitum*. For all experiments, behavioral testing was conducted during the light cycle.

### Drugs

The following drugs were used for behavioral pharmacology experiments: the α2AR agonist guanfacine hydrochloride (Sigma-Aldrich, St. Louis, MO), the peripheral aromatic acid decarboxylase inhibitor benserazide (Sigma-Aldrich), and the synthetic NE precursor l-3,4-dihydroxyphenylserine (DOPS; Lundbeck, Deerfield, IL). Guanfacine (0.3 mg/kg) was dissolved in sterile saline (0.9% NaCl) and injected i.p. at a volume of 10 ml/kg 30 min before behavioral testing. Sterile saline vehicle was injected to control for any confounding effect of injection stress on behavior, and vehicle-treated animals were used for statistical comparison.

For the DOPS rescue experiment, DOPS was dissolved in distilled water with 2% HCl, 2% NaOH, and 2 mg/kg vitamin C, as described (Schank et al. 2008; Thomas et al. 1998). *Dbh -/-* mice were injected s.c. with DOPS (1 g/kg) + benserazide (250 mg/kg), then tested 5 h later once NE levels peaked (Rommelfanger et al. 2007; Thomas et al. 1998). Benserazide was included to prevent the conversion of DOPS to NE in the periphery, thus restricting restoration of NE to the brain (Murchison et al. 2004). The vehicle control for the DOPS experiment was also administered 5 h before testing.

### Novelty-induced locomotion (NIL)

Individual mice were removed from their home cages and placed into large novel cages (10” × 18” × 10”) on locomotor activity testing racks (San Diego Instruments, San Diego, CA) (Schroeder et al. 2013; Weinshenker et al. 2002a). Novelty-induced ambulations (consecutive infrared beam breaks) were recorded for 1 h in 5 min bins. Testing occurred in a brightly lit room where the animals were housed. Test cages were covered with a lid and contained a thin layer of standard bedding substrate during NIL testing.

### Novelty-suppressed feeding (NSF)

All chow was removed from the home cage of subject animals 24 h prior to behavioral testing. On the day of testing, mice were moved to the test room and allowed to habituate for at least 2 h before starting the test. For pharmacological experiments, mice were injected either 30 min (guanfacine) or 5 h (DOPS) prior to testing. Individual mice were removed from their home cages and placed into a large novel cage (10” × 18” × 10”) with a single pellet of standard mouse chow in the center. The latency for the mouse to feed (grasp and bite the food pellet) was recorded using a stopwatch; mice that did not feed within 10 min were assigned a feeding latency score of 600 sec (Dulawa et al. 2005; Tillage et al. 2020). Testing occurred in a dark room under red light. Test cages were uncovered and contained a thin layer of standard bedding substrate.

### Feeding in the home cage and food neophobia

Mice were singly housed in clean standard mouse cages without chow for 24 h prior to behavioral testing to allow for cage familiarization. On the day of testing, mice were moved to the test room and allowed to habituate for at least 2 h before starting the test. For home cage feeding experiments, the cage lid was removed, and a standard food pellet was introduced into the cage. For food neophobia experiments, the cage lid was removed, and a standard food pellet coated in cinnamon (2 g cinnamon/100 g chow; Penzeys Spices, Wauwatosa, WI) was introduced into the cage (Modlinska and Stryjek 2016; Modlinska et al. 2015). The latency to feed was recorded for both tests using a stopwatch; mice that did not feed within 10 min were assigned a feeding latency score of 600 sec. Testing occurred in a dark room under red light.

### c-fos immunohistochemistry (IHC)

*Dbh -/-* and control mice were exposed to NIL or NSF tasks, after which they were left undisturbed in the test cage for 90 min (from the start of the task). The mice were then euthanized with an overdose of sodium pentobarbital (Fatal Plus, 150 mg/kg, i.p.; Med-Vet International, Mettawa, IL) for transcardial perfusion with cold 4% paraformaldehyde in 0.01 M PBS. Another group of *Dbh -/-* and control mice was selected for comparisons of brainwide c-fos induction under baseline, resting conditions; these animals were naïve to the behavioral tasks and were immediately perfused after removal from the home cage.

After extraction, brains were postfixed for 24 h in 4% paraformaldehyde at 4°C, and then transferred to cryoprotectant 30% sucrose/PBS solution for 72 h at 4°C. Brains were embedded in OCT medium (Tissue-Tek; Sakura, Torrance, CA) and serially sectioned by cryostat (Leica) into 40-µm coronal slices spanning the entire brain between the LC and orbitofrontal cortex. Brain sections were stored in 0.01 M PBS (0.02% sodium azide) at 4°C before IHC.

For IHC, brain sections were blocked for 1 h at room temperature in 5% normal goat serum (NGS; Vector Laboratories, Burlingame, CA) diluted in 0.01 M PBS/0.1% Triton-X permeabilization buffer. Sections were then incubated for 48 h at 4°C in NGS blocking/permeabilization buffer, including primary antibodies raised against c-fos (rabbit anti-c-fos, Millipore, Danvers, MA, ABE457; 1:5000) and the NE transporter (NET; chicken anti-NET, #260006, Synaptic System, Goettingen, Germany; 1:3000). After washing in 0.01 M PBS, sections were incubated for 2 h in blocking/permeabilization buffer with goat anti-rabbit AlexaFluor 488 and goat anti-chicken AlexaFluor 568 (Invitrogen, Carlsbad, CA; 1:500). After washing, the sections were mounted onto Superfrost Plus slides and coverslipped with Fluoromount-G plus DAPI (Southern Biotech, Birmingham, AL).

### Viral tracing of LC terminal fields in forebrain targets

Adult mice expressing Cre recombinase under the *tyrosine hydroxylase* promoter (TH-Cre; Jax #008601) were anesthetized with isoflurane (for induction and maintenance; Patterson Veterinary Supply, Devens, MA), placed in a stereotaxic frame (David Kopf Instruments, Tujunga, CA), and administered meloxicam (Patterson Veterinary Supply; 5 mg/kg, s.c.) for analgesia prior to the start of surgery. Mice received bilateral infusions of AAV10-EF1a-DIO-hChR2-eYFP (#20298, Addgene; Watertown, MA) into the LC (AP:−5.4 mm, ML: ±1.2 mm, DV, −4.0 mm) with a 5 µL Hamilton syringe (VWR, Radnor, PA) and stereotaxic injector pump (Stoelting, Wood Dale, IL). A total volume of 0.6 µL virus diluted 1:1 with sterile artificial cerebrospinal fluid (Harvard Apparatus, Holliston, MA) was infused per side at a rate of 0.15 µL/min. The infusion needle was left in place for 5 min after each infusion to allow for viral diffusion.

Mice were given at least 3 weeks to recover from surgery and allow full viral expression before they were euthanized and perfused, as described above. Brains were processed for IHC to visualize LC cell bodies and axon terminals in target regions using antibodies raised against TH (rabbit anti-TH, P40101-0, Pel-Freez, Rogers, AR; 1:1000) and YFP (chicken anti-GFP, ab13970, abcam, Cambridge, MA; 1:1000) All steps for IHC were identical to those described above except primary antibody incubations occurred over 24 h and primary antibodies were detected with goat anti-chicken 488 and goat anti-rabbit 568 (Invitrogen; 1:500).

### Fluorescent imaging and c-fos quantification

Fluorescent micrographs of immunostained sections were acquired on a Leica DM6000B epifluorescent upright microscope at 10x magnification with uniform exposure parameters for c-fos quantification. For the viral tracing experiment, micrographs were acquired at 20x for visualization of LC cell bodies and axon terminals. NET and TH antibodies were used to define LC cell bodies and identify noradrenergic axon terminals in target regions.

For c-fos quantification, we selected atlas-matched sections from each animal at the level of the LC and 15 of its forebrain targets. A standardized region of interest was drawn for all images to delineate the borders of discrete structures in all animals. Image processing and analysis were performed using ImageJ software. The analysis pipeline included background subtraction, intensity thresholding (Otsu method), and automated cell counting within defined regions of interest, guided by automated size and shape criteria for c-fos+ cells (size: 50–100 mm^2^, circularity: 0.6–1.0).

### Statistical analysis

For NIL experiments, the effect of time on locomotion in *Dbh -/-* and *Dbh +/-* mice was compared using a two-way repeated measures ANOVA (genotype x time), with post hoc Bonferroni’s tests for multiple comparisons. The within-trial habituation curves for NIL were fit with a simple linear regression for statistical comparison between *Dbh* genotypes. For NSF experiments, the effect of test cage familiarity on latency to feed after food deprivation was compared by two-way ANOVA (genotype x test environment), with post hoc Sidak’s tests for multiple comparisons. Food neophobia between genotypes, NSF behavior between DOPS- and vehicle-treated *Dbh -/-* mice, and NSF behavior between guanfacine- and vehicle-treated *Dbh +/-* control mice were analyzed using unpaired t-tests.

For c-fos quantification, genotype differences were compared in the LC and 15 forebrain targets at baseline and after NIL or NSF. Comparisons were made within behavioral tasks and between genotypes by multiple t-tests using the Holm-Sidak correction for multiple comparisons. The threshold for adjusted significance was set at *p* < 0.05, and two-tailed variants of tests were used throughout. Graphical data are presented as group mean ± SEM. Analyses and graph design were performed using Prism v8 (GraphPad Software, San Diego, CA).

## Results

### Norepinephrine deficiency attenuates novelty-induced locomotion and promotes rapid habituation

To determine the contribution of NE to neophilic behavior, a cohort of age- and sex-matched *Dbh -/-* and *Dbh +/-* control mice were compared in the NIL test, with locomotor activity measured in 5 min bins across 1 h. A two-way repeated measures ANOVA (genotype x time) showed a main effect of genotype (F(1,21) = 9.28, *p* < 0.01), time (F(11,231) = 30.16, *p* < 0.0001), and a time x genotype interaction (F(11,231) = 2.81, *p* = 0.02). *Dbh -/-* mice made fewer novelty-induced ambulations overall compared to control mice (1612±582 vs 2405±659), and post hoc analyses revealed that the *Dbh -/-* mice were less active than control mice at the beginning of the task during the 5 min (*p* = 0.01), 10 min (*p* < 0.01), 15 min (*p* = 0.03), and 20 min (*p* = 0.03) time bins (Fig. 1a).

**Fig. 1.**
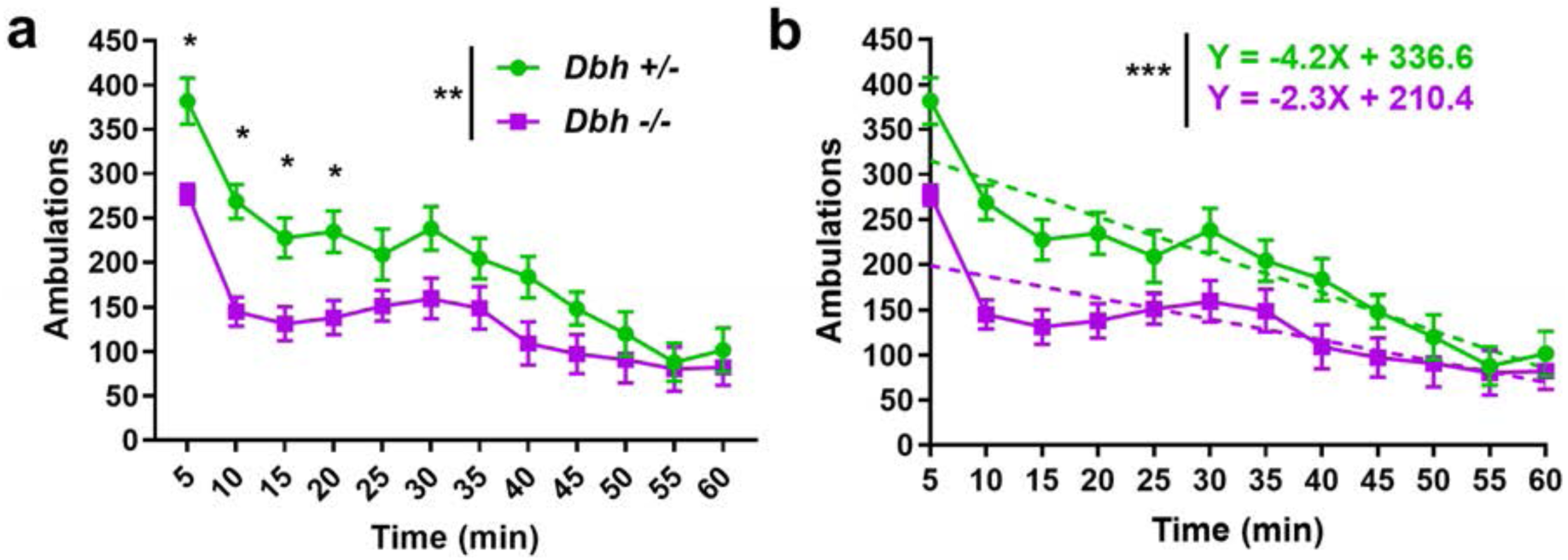
Assessment of neophilia in *Dbh -/-* and *Dbh +/-* mice. **a** Novelty-induced locomotor behavior was attenuated in *Dbh -/-* mice (n = 11) compared to controls (n = 12). Mice of both *Dbh* genotypes demonstrated the most activity at the start of the task followed by habituation, with activity diminished over time. However, in the four earliest time bins of the test, *Dbh -/-* mice were less active than controls. **b** The change in novelty-induced locomotion over time for both *Dbh* genotypes was assessed using a simple linear regression analysis. The slope of the dashed fit lines of within-trial habituation to the novel environment indicates a blunted initial response to novelty in *Dbh -/-* mice (represented by the Y-intercept) and a flatter slope, suggesting more rapid habituation to the novel environment. Error bars ± SEM, \*\*\**p* < 0.001, **p *<* 0.01, *p < 0.05.

Mice of both genotypes showed functional novelty detection, exhibiting the most activity in time bins at the beginning of the task and within-trial habituation to the novel environment. However, simple linear regression analysis of ambulations over time revealed that variation in locomotor activity was predicted more strongly by variation in time in the control mice (*R*^*2*^ *=* 0.43) than in *Dbh -/-* mice (*R*^*2*^ *=* 0.23). There was a significant difference in the slope of the fit lines for each *Dbh* genotype (F(1,272) = 11.22, *p* < 0.001). The flatter slope of the *Dbh -/-* fit line suggests more rapid locomotor habituation to the novel environment in NE-deficient mice (Fig. 1b).

### Central norepinephrine is necessary and sufficient for neophobia in the novelty-suppressed feeding test

To assess whether NE is also required for neophobia, the behavior of *Dbh -/-* and control mice were compared in the NSF test. A two-way ANOVA (genotype x environment) showed a main effect of genotype (F(1,48) = 32.62, *p* < 0.0001), environment (F(1,48) = 30.38, *p* < 0.0001), and a genotype x environment interaction (F(1,48) = 25.39, *p* < 0.0001). Post hoc comparisons revealed that in the novel test environment, *Dbh -/-* mice ate much more rapidly than control mice (91.93±44.43 vs 389.49±162.81 s, *p* < 0.0001), but did not differ from control mice when assessed for feeding latency in the home cage (78.82±44.79 vs 92.45±47.43 sec, *p* > 0.05) (Fig. 2a). Thus, the genotype differences in feeding latency are likely to be related to reduced neophobia rather than increased hunger. As expected, *Dbh +/-* mice ate significantly more rapidly in the home cage than they did in the novel cage (t(48) = 7.57, *p* < 0.0001). However, latency to feed did not differ between test environments in *Dbh -/-* mice (t(48) = 0.14, *p* > 0.05), suggesting NE deficient mice exhibit behavioral indifference to context familiarity when hunger is a motivating factor.

**Fig. 2.**
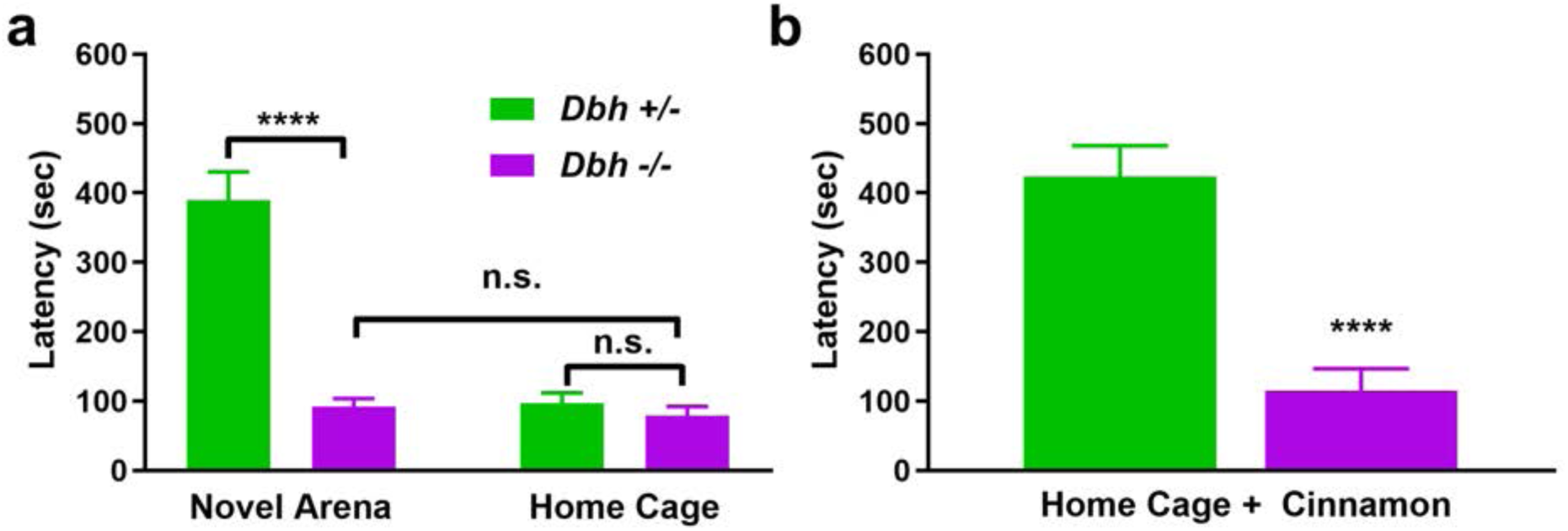
Assessment of novelty-suppressed feeding and food neophobia in *Dbh -/-* and *Dbh +/-* mice. **a** Compared to *Dbh* +/-controls (n = 16), *Dbh -/-* (n = 14) ate familiar mouse chow much more rapidly in a novel environment. Unlike controls, *Dbh -/-* mice did not have a significant difference in feeding latency between the novel test environment and familiar home cage. The latency to feed did not differ between *Dbh* +/-controls (n = 11) and *Dbh -/-* mice (n = 11) when tested in the home cage. **b** Compared to *Dbh* +/-controls (n = 11), *Dbh -/-* mice (n = 10) ate the novel-scented cinnamon chow more rapidly in a familiar environment, indicating diminished food neophobia. Error bars ± SEM, \*\*\*\**p* < 0.0001, n.s. = not significant.

We also measured the effects of NE deficiency in a variation of the NSF test with novel-scented food presented in a familiar environment. To this end, we coated standard mouse chow pellets with cinnamon, a novel olfactory stimulus that is neither innately appetizing nor aversive to rodents (Modlinska and Stryjek 2016; Modlinska et al. 2015; Scholtysik 1980). Similar to the novel environment version of the test, *Dbh* -/- mice ate the novel-smelling pellet much more rapidly than control mice (115.3±99.7 vs 423.6±148.1 sec; t(19) = 5.54, *p* < 0.0001) (Fig. 2b).

To evaluate whether replacing central NE in *Dbh -/-* mice would restore neophobic behavior in *Dbh -/-* mice, we compared knockouts treated with vehicle or the synthetic NE precursor DOPS in the NSF test. DOPS treatment significantly increased the latency to feed in *Dbh -/-* mice compared to vehicle treatment (366.4±99.3 vs 88.8±48.8 sec; t(9) = 6.07, *p* < 0.001), rescuing feeding latency to similar levels as control mice (Fig. 3a). To determine whether acutely disrupting NE transmission in normal mice confers resistance to neophobia, we also compared *Dbh +/-* mice treated with vehicle or guanfacine (0.3 mg/kg) in the NSF test. Guanfacine is an agonist of α2a inhibitory autoreceptors and inhibits NE release (Fox et al. 2012). Guanfacine produced a knockout-like phenotype in *Dbh +/-* mice, significantly decreasing latency to feed in the novel environment (312.1±56.1 vs 525.3±76.4 sec; t(16) = 6.58, *p* < 0.0001) (Fig. 3b). Thus, neophobia can be restored in NE deficient mice by pharmacological rescue of brain NE synthesis, and reduced in control mice with the anti-adrenergic drug guanfacine.

**Fig. 3.**
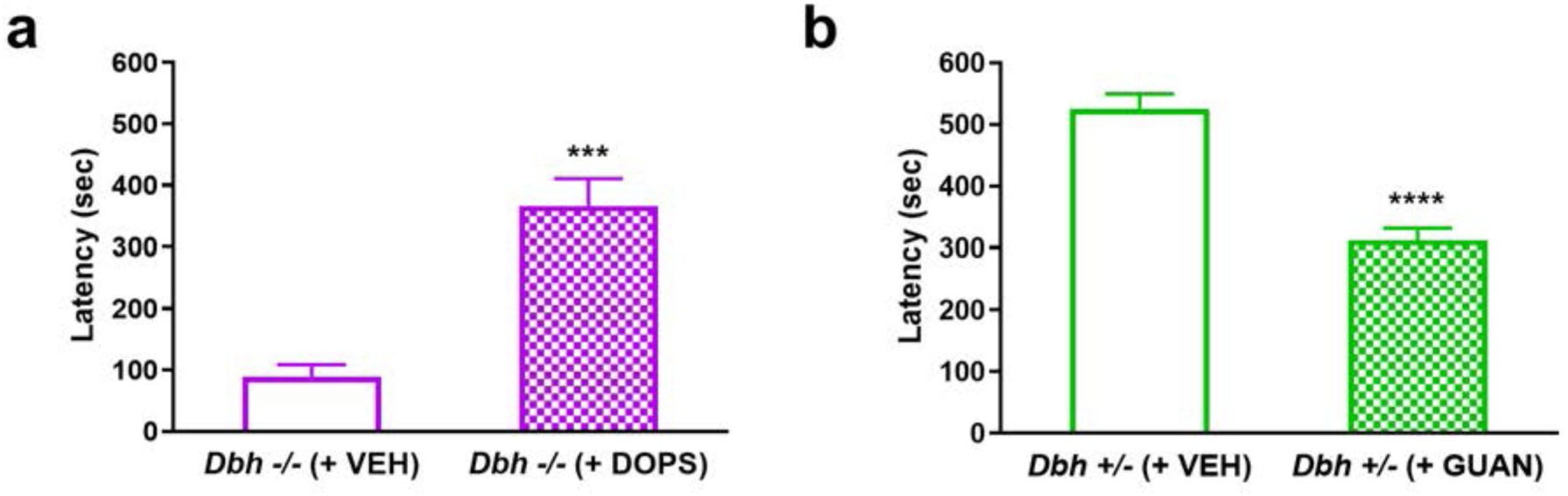
Pharmacological restoration and attenuation of neophobia. **a** *Dbh -/-* mice in which central NE was restored with DOPS (n = 5) before NSF testing demonstrate significantly longer latencies to feed than *Dbh -/-* mice treated with vehicle (VEH; n = 6). **b** *Dbh +/-* control mice in which central NE transmission was reduced with guanfacine (n = 8) ate more rapidly than control mice treated with VEH (n = 10) in the NSF test. Error bars ± SEM, \*\*\*\**p* < 0.0001, \*\*\**p* < 0.001.

### LC neurons project to forebrain regions implicated in responses to novelty

To determine whether LC neurons project to forebrain region implicated in novelty responses, we bilaterally injected a viral vector encoding Cre-dependent ChR2-eYFP into the LC of TH*-*Cre mice to label noradrenergic LC neurons and their and their axons and axon terminals (Fig. 4a). Brainstem cell body expression of eYFP was histologically confirmed for all cases and was restricted to TH+ neurons in the LC (Fig. 4b).

**Fig. 4.**
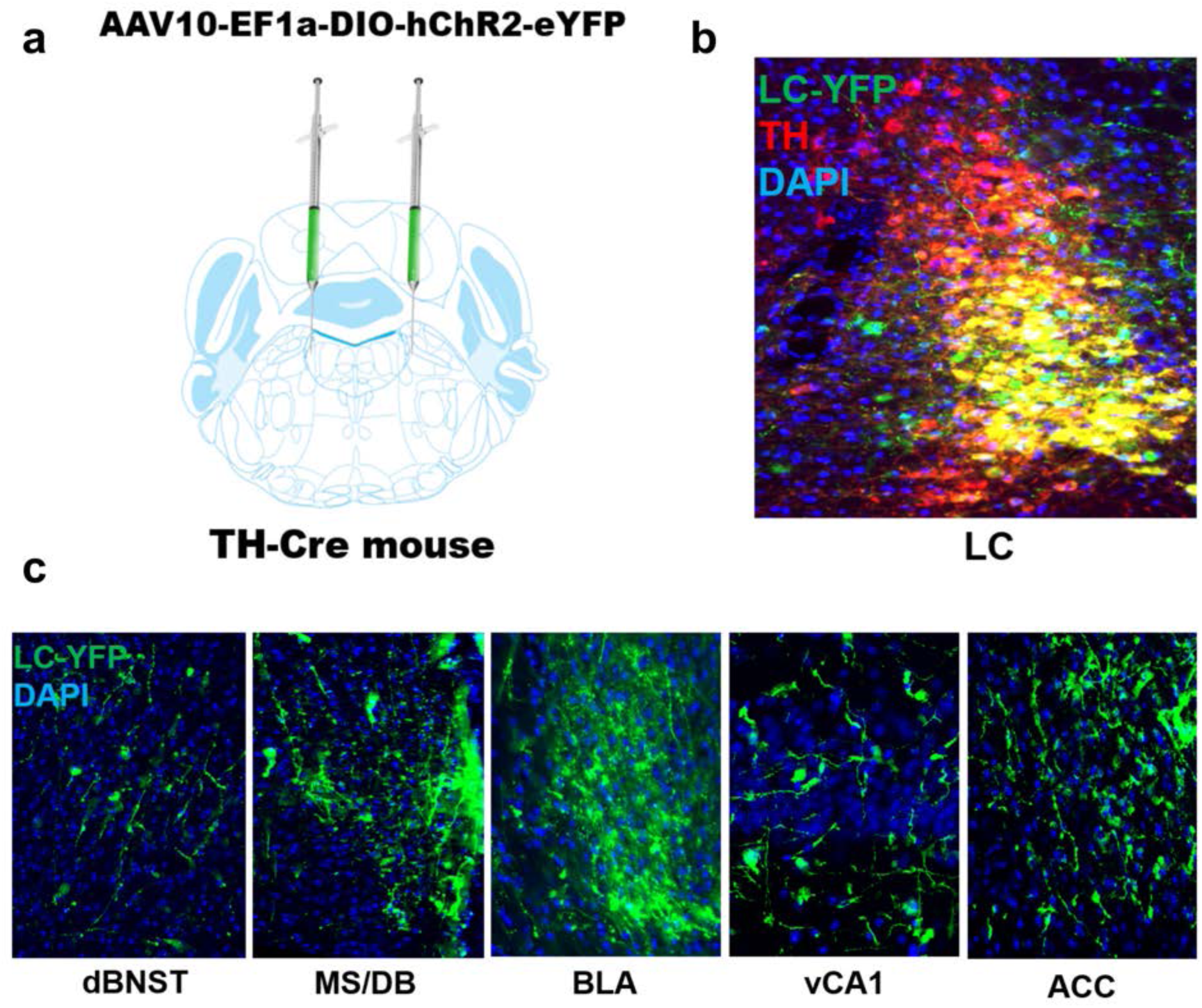
Viral tracing of locus coeruleus (LC) fibers in forebrain regions implicated in novelty. **a** TH-Cre mice received bilateral intra-LC infusions of a Cre-dependent viral vector encoding ChR2-eYFP, which becomes concentrated in axons and axon terminals and can be used as a tool for projection mapping. **b** ChR2-YFP expression (green) substantially overlaps with immunostaining for TH (red) in the LC and is largely restricted to TH+ neurons. **c** Representative micrographs of eYFP+ terminals from the LC in the dorsal bed nucleus of the stria terminals (dBNST), medial septum/diagonal band complex (MS/DB), basolateral amygdala (BLA), the CA1 subfield of the ventral hippocampus (vCA1), and the anterior cingulate cortex (ACC).

We next examined 15 forebrain regions previously implicated in novelty responses and reported to receive noradrenergic innervation from the LC (Robertson et al. 2016; Schwarz and Luo 2015; Uematsu et al. 2015; Zaborszky 1989), including the medial orbitofrontal cortex (mOFC), prelimbic (PrL) and infralimbic cortices (IL), anterior cingulate cortex (ACC), agranular insula (AI), claustrum (CL), medial septum/diagonal band (MS/DB), dorsal bed nucleus of the stria terminalis (dBNST), basolateral amygdala (BLA), dorsal dentate gyrus (dDG), dorsal hippocampal subfields CA1 (dCA1) and CA3 (dCA3), ventral hippocampus subfield CA1 (vCA1), lateral hypothalamus (LH), and the paraventricular nucleus of the thalamus (PVT). All forebrain regions analyzed contained eYFP+ fibers, demonstrating direct inputs from the LC to novelty-sensitive forebrain targets (Fig. 4c).

### Norepinephrine deficiency is associated with blunted c-fos induction in several forebrain targets of the locus coeruleus following novelty exposure

At baseline, c-fos expression was negligible in all regions examined, and there were no significant differences in c-fos+ cells in any region between *Dbh* genotypes (*p* > 0.05 for all regions examined, data not shown). To identify potential NE-sensitive novelty circuits from among the identified forebrain LC targets, *Dbh -/-* and *Dbh +/-* control mice were euthanized for quantification of c-fos+ cells in the LC and 15 the brain regions (Fig. 5a) identified following either NIL or NSF.

**Fig. 5.**
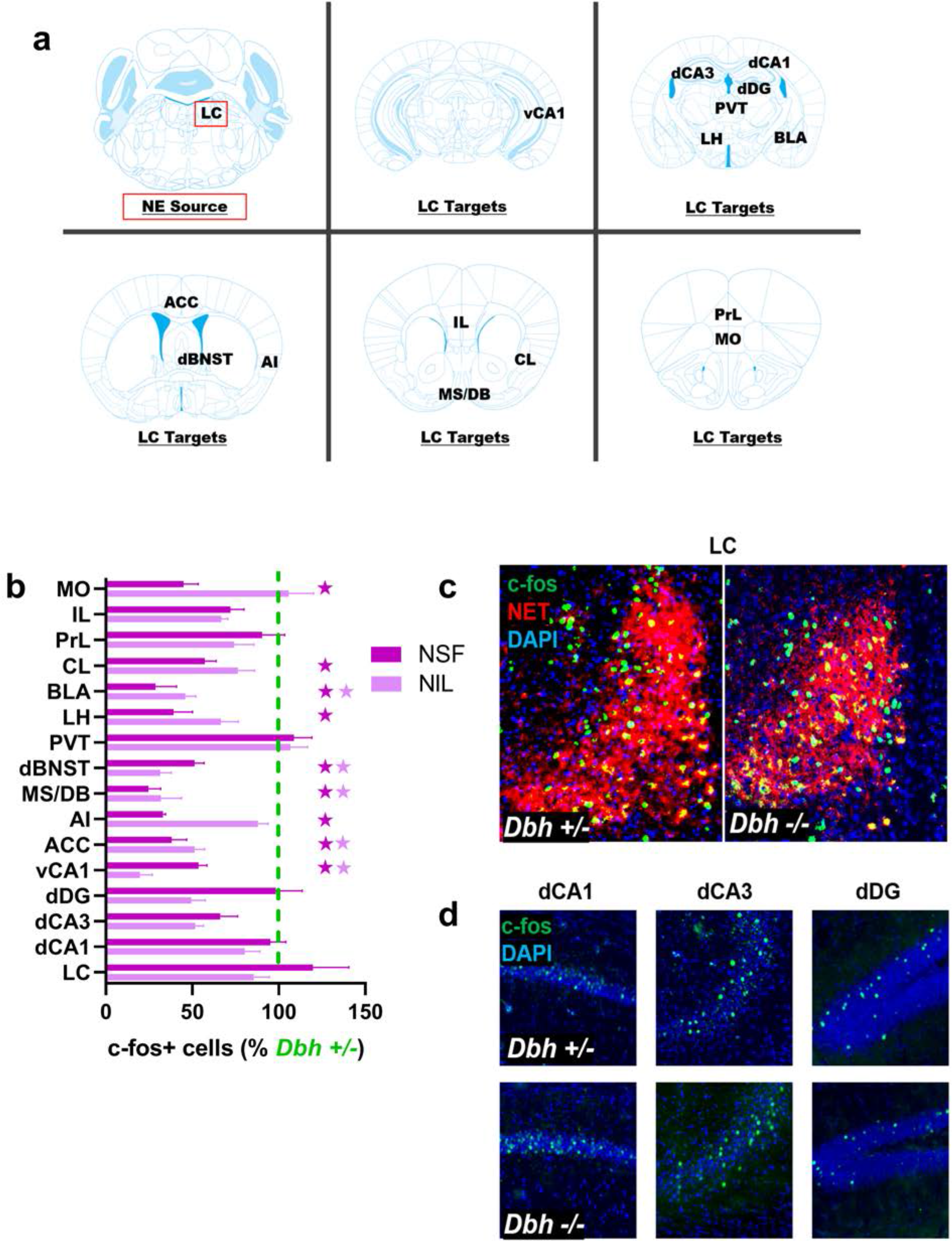
Assessing the effects of NE deficiency on neuronal activity in the locus coeruleus (LC) and forebrain target regions. **a** Anatomical location of the LC and 15 target regions in the forebrain: medial orbitofrontal cortex (MO), prelimbic (PrL) and infralimbic cortices (IL), anterior cingulate cortex (ACC), agranular insula (AI), claustrum (CL), medial septum/diagonal band (MS/DB), dorsal bed nucleus of the stria terminalis (dBNST), basolateral amygdala (BLA), dorsal dentate gyrus (dDG), dorsal hippocampal subfields CA1 (dCA1) and CA3 (dCA3), ventral hippocampus subfield CA1 (vCA1), lateral hypothalamus (LH), and the paraventricular nucleus of the thalamus (PVT). **b** c-fos induction in *Dbh -/-* mice in the LC and forebrain targets after NSF (darker purple) and NIL (lighter purple), expressed as averaged percentage of c-fos+ cell counts in *Dbh +/-* controls for each test. Dark purple stars indicate *p <* 0.05 for NSF, light purple starts indicate *p <* 0.05 for NIL. **c** Representative micrographs showing similar levels of c-fos (green) induction between *Dbh* genotypes after NSF in NET+ LC neurons (red). **d** *Dbh* genotype does not affect the number of c-fos+ cells observed following NSF in the dorsal hippocampus.

Exposure to the novel environment in either test resulted in robust c-fos induction in the LC regardless of *Dbh* genotype (t(9) = 0.69, *p* > 0.05 for NIL; t(6) = 0.81, *p* > 0.05 for NSF) (Fig. 5b,c), suggesting that NE deficiency does not impair LC activation in response to spatial novelty. c-fos induction (compared to baseline) was observed following NIL or NSF tests in LC target regions including the PVT, PrL, IL, dCA1, dCA3, and dDG, but the number of c-fos+ cells did not differ between genotypes (*p* > 0.05 for all comparison) (Fig. 5b,d).

By contrast, profound genotype differences in c-fos induction emerged in several other forebrain targets of the LC during following NIL and NSF. Following either behavioral tasks, *Dbh -/-* mice had significantly fewer c-fos+ cells than control mice in vCA1 (t(9) = 5.02, *p* < 0.01 for NIL; t(9) = 5.13, *p* < 0.01 for NSF), ACC (t(9) = 4.86, *p* = 0.01 for NIL; t(9) = 5.20, *p* < 0.01 for NSF), dBNST (t(8) = 9.12, *p* < 0.001 for NIL; t(9) = 4.24, *p* = 0.02 for NSF), MS/DB (t(8) = 5.25, *p* < 0.01 for NIL; t(8) = 6.63, *p* < 0.01 for NSF), and BLA (t(8) = 5.92, *p* < 0.01 for NIL; t(9) = 5.63, *p* < 0.01 for NSF) (Fig.5b; Fig. 6a). After NSF, but not after NIL, NE-deficient mice also had fewer c-fos+ cells than controls in MO (t(9) = 5.63, *p* < 0.01), CL (t(9) = 4.38, *p* = 0.02), AI (t(9) = 3.69, *p* = 0.04), and LH (t(9) = 5.69, *p* < 0.01) (Fig. 5b; Fig. 6b).

**Fig. 6.**
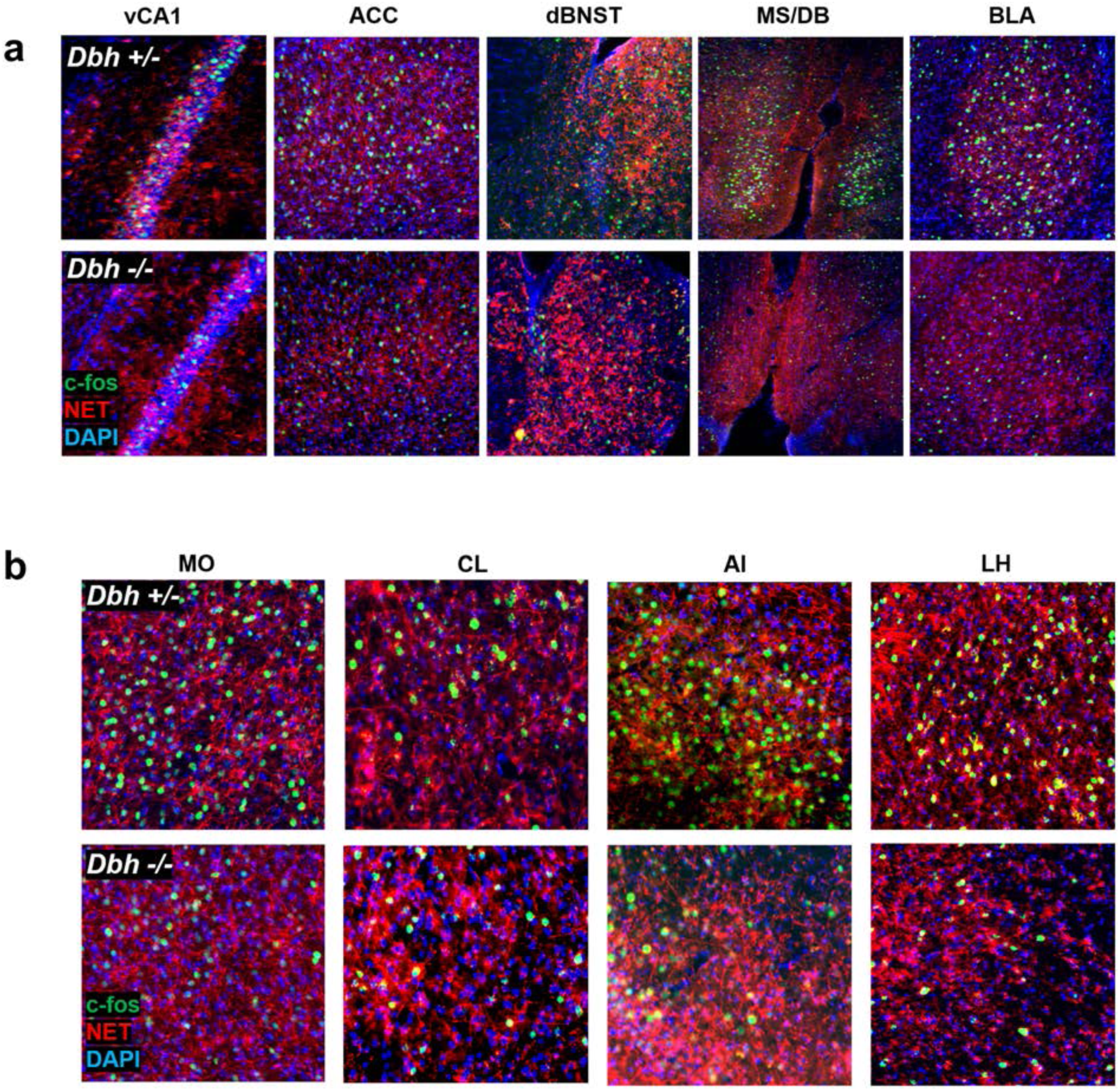
Visualization of NET+ noradrenergic fibers and c-fos induction in target regions after novelty-suppressed feeding (NSF). **a** Representative micrographs showing immunostaining for NET (red) and c-fos (green) between *Dbh* genotypes after NSF. Following either NSF or NIL, *Dbh -/-* mice showed fewer c-fos+ cells than controls in ventral hippocampus subfield CA1 (vCA1), anterior cingulate cortex (ACC), dorsal bed nucleus of the stria terminalis (dBNST), medial septum/diagonal band (MS/DB), and basolateral amygdala (BLA). **b** Following NSF, but not following NIL, *Dbh -/-* mice showed fewer c-fos+ cells than controls in medial orbitofrontal cortex (MO), claustrum (CL), agranular insula (AI), and lateral hypothalamus (LH).

## Discussion

### Norepinephrine is required for sustained locomotor activation in novel environments

In this study, we first compared neophilic behavior in NE-deficient and NE-competent littermate control mice in the NIL test. We found that over the course of 1 h, NE-deficient mice exhibited reduced locomotor activity and more rapid habituation in the novel test environment compared to control mice. These findings are congruent with previous reports from our lab and suggest that novelty may have reduced incentive value or intrinsic salience in NE-deficient animals (Cubells et al. 2016; Porter-Stransky et al. 2019; Weinshenker et al. 2002a).

Importantly, novel environments elicit an initial increase in locomotion in *Dbh -/-* mice that habituates over time, suggesting that spatial memory mechanisms that distinguish novel and familiar environments remain intact in the absence of NE. Previous work has also demonstrated functional single-trial learning of a novel environment in *Dbh -/-* mice; total locomotion during the first exposure to the novel environment was significantly greater than total locomotion during re-exposure to the same environment 24 h later in both *Dbh -/-* and control mice (Weinshenker et al. 2002a). It is interesting to note that this particular type of “one-shot” contextual learning has recently been shown to depend on LC transmission of dopamine (DA) to the dorsal hippocampus (Kempadoo et al. 2016; Takeuchi et al. 2016; Wagatsuma et al. 2018). Because DBH catalyzes the conversion of DA to NE, *Dbh -/-* mice make DA instead of NE in LC neurons, and thus LC-DA transmission would theoretically be undisturbed (or even augmented) in *Dbh -/-* mice (Bourdélat-Parks et al. 2005; Schank et al. 2006; Weinshenker et al. 2002b).

### Norepinephrine is necessary and sufficient for novelty-suppressed feeding

The present study also investigated the effect of NE deficiency on neophobia in the NSF test. NE-deficient mice demonstrated a total absence of neophobia in the novel test environment that could be rescued to control levels by restoring central NE synthesis with DOPS. In control mice, the anti-adrenergic drug guanfacine produced rapid anxiolytic effects when administered prior to the NSF test. Together, these findings demonstrate that NE is both necessary and sufficient for species-typical neophobia in NSF. Importantly, feeding latencies in NE-deficient mice did not differ from controls in the familiar home cage environment, ruling out the possibility that NE-deficient mice are simply hungrier after food deprivation than NE-competent mice (Cryan and Sweeney 2011; Dulawa et al. 2005).

The profound lack of anxiety displayed by the *Dbh -/-* mice in the NSF test contrasts sharply with the seemingly “normal” phenotype of these mice in canonical tests of anxiety, including the elevated plus and zero mazes (EPM/EZM), light/dark box (LDB), and open field test (OFT) (Lustberg et al. 2019; Marino et al. 2005; Schank et al. 2008). These conflict-based models of anxiety-like behavior are widely used in rodent studies, but they rely on the underlying assumption that mice are motivated to explore novel environments (open arms of EPM/EZM, light compartment of LDB, center of OFT). The conflict in these canonical tasks is between innate fear (open arms, light, center of field) and the drive to explore the novel environment (Cryan and Sweeney 2011; Kalueff et al. 2007). Given that *Dbh -/-* mice exhibit blunted exploratory behavior in novel environments, it is possible that any anxiolytic phenotype would be occluded by a lack of neophilia. In other words, the behavioral measures of anxiolysis in these canonical anxiety tasks is exploratory behavior, which in NE-deficient mice is confounded by attenuated neophilia. In NSF, the conflict occurs between the fear of the novel environment and the motivation to eat. The behavioral measure of anxiolysis in NSF is latency to feed; thus, curiosity and neophilia are not a factor in this task.

The fact that a robust anxiolytic phenotype emerged in *Dbh -/-* mice during NSF but not in more commonly employed tests of anxiety-like behavior should serve as a reminder that any behavioral model has underlying assumptions and associated confounds (Cryan and Sweeney 2011; Kalueff et al. 2007). We urge researchers investigating anxiety-like behavior to include NSF in the battery of behavioral testing in order to account for potentially confounding effects of neophilia in more canonical anxiety tests, which all fundamentally measure exploration.

### NE-deficient mice show reduced c-fos induction in select targets of the LC following NIL and NSF

Following exposure to the novel environment in the NIL and NSF tests, robust c-fos induction was observed in both NE-deficient and control LC. This finding suggests that LC activation in response to novelty does not depend on NE signaling and may be induced by glutamatergic inputs to the LC, or an excitatory neuropeptide transmitter such as orexin or corticotropin-releasing factor (CRF) (Gompf et al. 2010; Soya et al. 2017; Valentino et al. 1993).

Despite demonstrating similar levels of LC activation compared to controls after NIL or NSF, *Dbh -/-* mice had fewer c-fos+ cells in several forebrain targets of the LC, including vCA1, ACC, dBNST, MS/DB, and BLA. Because these regions were hypoactive in the absence of NE following novelty exposure in both tests, we propose that they comprise the nodes of a noradrenergic novelty network (NNN) that drives both neophilic and neophobic behavior (Fig. 7). With outputs extending throughout the prefrontal cortex, striatum, hypothalamus, midbrain, and brainstem, the NNN is well positioned to coordinate affective and motor responses to unfamiliar environments (Lebow and Chen 2016; Li and Wang 2018; Padilla-Coreano et al. 2016; Stachniak et al. 2014; Tovote et al. 2015).

**Fig. 7.**
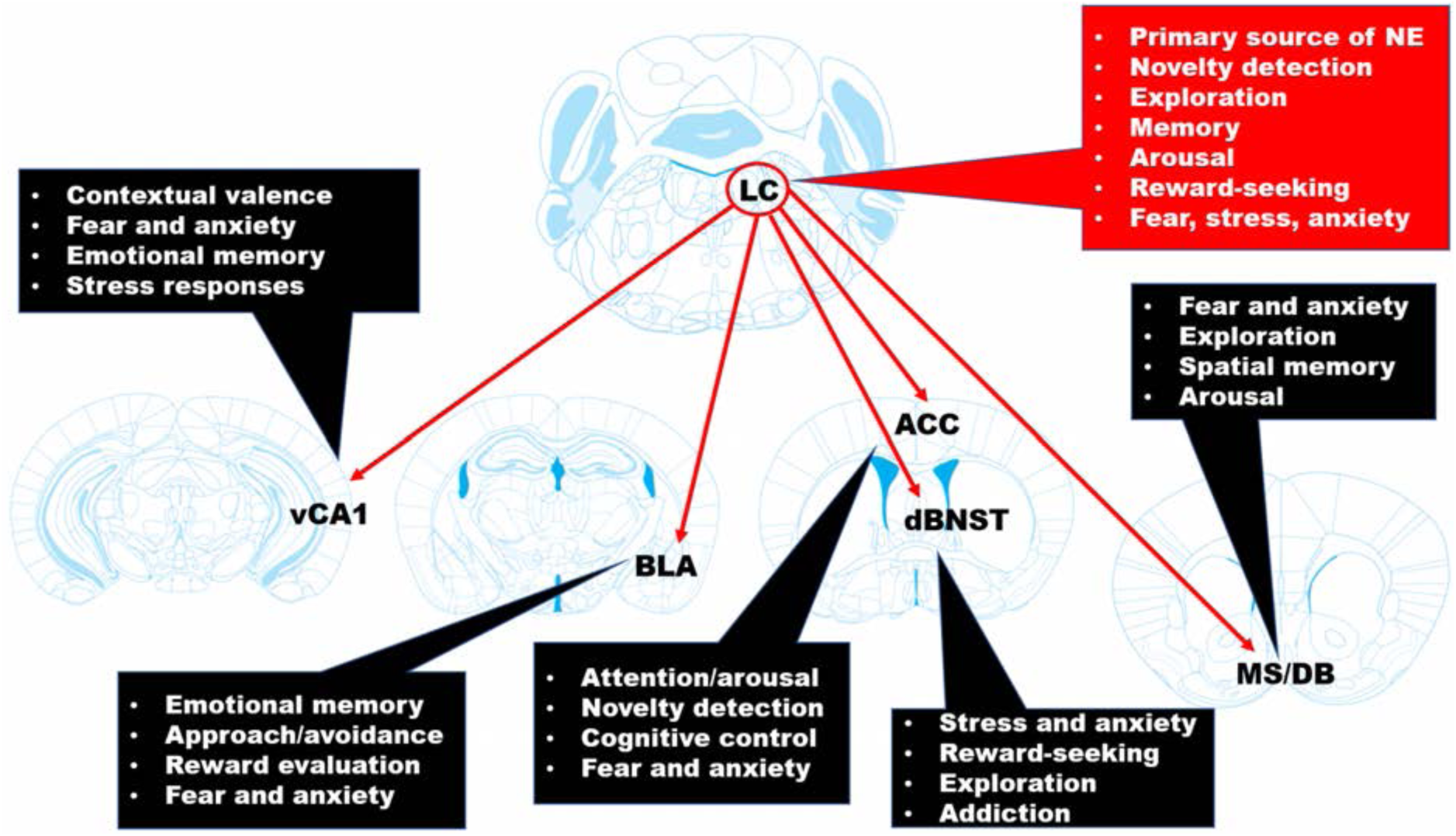
The noradrenergic novelty network (NNN). LC-NE inputs to ventral hippocampus subfield CA1 (vCA1), anterior cingulate cortex (ACC), dorsal bed nucleus of the stria terminalis (dBNST), medial septum/diagonal band (MS/DB), and basolateral amygdala (BLA) may support diverse and flexible behavioral responses in novel environments. The outputs of the various nodes of the NNN could “prime” either fear-related or exploratory-related circuits in unfamiliar contexts, thus controlling both neophobic and neophilic behavior. Dysregulation of the NNN could be a common feature in certain psychiatric conditions, including anxiety and substance abuse disorders.

Recently, a population of spatially- and emotionally-tuned cells in vCA1 were shown to fire preferentially in anxiogenic environments (Jimenez et al. 2018). These “anxiety cells” are analogous to place cells in dCA1 which support spatial learning and memory, and exert their anxiogenic behavioral effects through excitatory glutamatergic projections to LH (Jimenez et al. 2018). The ventral hippocampus has been implicated in emotional learning and neophobia (Bannerman et al. 2002; Bannerman et al. 2004; Fanselow and Dong 2010; Padilla-Coreano et al. 2016; Santarelli et al. 2003), but the effects of manipulating LC-NE signaling within vCA1 on neophobia have not been described.

Reciprocal connections between the LC and the ACC support vigilance in novel environments, and destruction of LC terminals in the ACC suppresses electrophysiological measures of arousal associated with exposure to contextual novelty (Gompf et al. 2010). Recently, our group reported hypoactivity in the ACC of NE-deficient mice following cage change stress or exposure to a novel environment containing marbles (Lustberg et al. 2019). Unlike control mice, NE-deficient mice demonstrated virtually no stress-induced repetitive behaviors following cage change or novelty exposure, suggesting that NE engagement of the ACC may be necessary for typical affective responses to contextual change (Egner 2011).

The dBNST is a region with a well-established role in innate fear, stress responses, and anxiety-like behavior (Avery et al. 2016; Lebow and Chen 2016). CRF-containing neurons in the dBNST are activated by NE and engage the hypothalamic-pituitary-adrenal axis to synchronize endocrine and behavioral responses to stress (Dabrowska et al. 2016; Egli et al. 2005; Vranjkovic et al. 2017). Activation of this region is also implicated in stress-induced reinstatement of drug-seeking behavior (Mantsch et al. 2016), an animal model of relapse, as well as threat-detection and stimulus evaluation under conditions of uncertainty (Lebow and Chen 2016).

Cholinergic neurons in the MS/DB of the basal forebrain are densely innervated by the LC (Bergado et al. 2007; Schwarz and Luo 2015; Schwarz et al. 2015), and control both exploratory and anxious behavior in novel environments (Bannerman et al. 2004; Carpenter et al. 2017; Myhrer 1989; Zhang et al. 2017). These cholinergic projection neurons likely modulate exploratory and avoidant behavior in novel environments via projections to the hippocampus and medial prefrontal cortex (Jiang et al. 2018; Tereshchenko et al. 2008). Although LC inputs to the MS/DB have not been well characterized, pharmacological studies suggest that the NE increases excitability of cholinergic neurons in this region (Berridge and Espana 2000; Berridge et al. 1996).

The BLA is critical for fear learning and anxiety-like behavior, but is also implicated in novelty responses (Balderston et al. 2011; Jhang et al. 2018). LC-NE signaling within the BLA elicits anxiety-like behavior in canonical anxiety tests as well measures of social anxiety (McCall et al. 2017; Siuda et al. 2016), but the role of the LC-NE → BLA circuit in neophobia has not been described.

### NE-deficient mice show diminished c-fos induction in select targets of the LC following NSF

After NSF, but not after NIL, NE-deficient mice had fewer c-fos+ cells than controls in MO, AI, CL, and LH. The MO is a behavioral control center at the anterior pole of the brain that is bidirectionally connected to the LC (Aston-Jones and Cohen 2005; Rolls and Grabenhorst 2008; Sadacca et al. 2017). MO neurons participate in the evaluation of risk and reward, decision making, and executive control of emotional responses (Petrides 2007; Woon et al. 2019). The AI is a cortical region involved in processing gustatory and interoceptive information (Naqvi and Bechara 2010; Uddin et al. 2017), and it is implicated in panic and anxiety disorders (Gehrlach et al. 2019; Klumpp et al. 2012; Poletti et al. 2015; Wittmann et al. 2014). NE signaling within the AI mediates food neophobia and is derived from the LC as well as non-cerulean brainstem groups (Robertson et al. 2016; Rojas et al. 2015). The CL is located between the AI and striatum, has connectivity with virtually all cortical regions, and is involved in contextual memory, novelty detection, allocation of attention, and action selection (Brown et al. 2017; Crick and Koch 2005; Qadir et al. 2018).

The LH is comprised of a heterogenous population of neurons that are also bidirectionally connected to the LC and participate in feeding, anxiety, stress responses, and motivated behavior (Bonnavion et al. 2016; Tanaka et al. 2000). One possible reason that NE-deficient mice demonstrated hypoactivity in these regions after NSF but not after NIL may be the relative demands and parameters of these two tests. For instance, NSF requires the animals to be food-deprived before testing so that the mice are motivated to eat in the test. On test day, NSF also requires the animals to make a behavioral “choice” between avoidance and approach to the food pellet in the novel environment. Thus, NE transmission in NSF may modulate gustatory reward-evaluation and action selection circuits in MO, AI, and CL, as well as motivational and emotional circuits within the LH in the context of the NSF test. The NIL test does not require food deprivation or decision making, potentially masking genotype differences in NE-dependent c-fos induction in these regions.

### NE-deficient mice show similar c-fos induction to controls in dorsal hippocampus and paraventricular thalamus after following NIL and NSF

By definition, novelty detection requires engagement of memory systems (Kafkas and Montaldi 2018). In order to determine that an environment is novel, it must be compared with existing representations of previously encountered environments that are stored in memory. Despite their striking absence of neophilia and neophobia, NE-deficient mice retain the ability to recognize an environment as novel or familiar (Weinshenker et al. 2002a). The dissociation between novelty detection and novelty responding in the NE-deficient animals suggests at least partially non-overlapping neural substrates for these operations (Bannerman et al. 2002; Harro et al. 1995; Kafkas and Montaldi 2014; Tereshchenko et al. 2008; Wingo et al. 2016).

Studies in rodents suggest that the hippocampus is functionally segregated along the dorsoventral axis; the dorsal hippocampus is required for spatial memory and navigation, while the ventral hippocampus is required for emotional memory and anxiety (Bannerman et al. 2002; Fanselow and Dong 2010). Although NE-deficient mice had reduced emotional responses and diminished neuronal activity in vCA1 after novelty exposure, they showed no impairment in the detection of spatial novelty and displayed neuronal activity similar to controls in dCA1, dCA3, and dDG after novelty exposure. *Dbh -/-* mice also had similar levels of activity compared to controls in the PVT, a midline thalamic structure that integrates complex contextual signals from hypothalamus and brainstem and which becomes highly activated under conditions of uncertainty (Choi and McNally 2017; Kirouac 2015). As mentioned above, LC-DA transmission is theoretically intact in NE-deficient mice, and recent studies have shown that LC-DA transmission excites neurons within the dorsal hippocampus and PVT (Beas et al. 2018; Wagatsuma et al. 2018).

### Limitations and future directions

A limitation of our study is that we cannot infer a causal role for LC-NE transmission to any of the identified forebrain regions in the expression of NIL and NSF behaviors. Additional experiments using optogenetic and chemogenetic tools to bidirectionally manipulate distinct forebrain targets of the LC are necessary to functionally dissect the NNN and understand how NE organizes affective responses to novelty.

Although we have clearly demonstrated a critical role for central NE transmission in neophobia, the neurochemistry underlying innate anxiety is complex and likely to involve multiple neuromodulatory systems. Intriguingly, serotonin (5-HT) deficient mice (*Tph2 -/-*) also exhibit dramatic reductions in neophobia in the NSF test (Angoa-Pérez et al. 2012; Mosienko et al. 2012). Moreover, these 5-HT deficient mice display increased aggressive and compulsive behaviors, which are absent in NE deficient mice (Angoa-Pérez et al. 2012; Lustberg et al. 2019; Marino et al. 2005; Mosienko et al. 2012). Indeed, the phenotypes of *Tph2 -/-* and *Dbh -/-* mice are perfectly opposed with the notable exception of reduced neophobia in both mutants, suggesting that behavioral expression of neophobia may require both NE and 5-HT (Blier and El Mansari 2007).

## Conclusions

In summary, these findings support an expanded role of central NE transmission in the expression of neophilic and neophobic behaviors. Guanfacine and related drugs have been used both on- and off-label for the treatment of anxiety and substance abuse disorders (Fox et al. 2012; Hoehn-Saric et al. 1981), but this is the first study to demonstrate anxiolytic effects of guanfacine in the NSF test. We propose that the anti-adrenergic agents like guanfacine should be investigated for rapid anxiolytic effects in patients with context-specific anxiety symptoms, such as agoraphobia (Marazziti et al. 2012; Wittmann et al. 2014).

## Acknowledgments

We thank Lundbeck for providing the DOPS. This work was supported by the National Institutes of Health (AG061175, NS102306, and DA038453 to DW; GM8602-22 to DL; MH116622 to RPT).

